# Mutation E300 recommended by protein language models gives ChrimsonR amplified photocurrent response

**DOI:** 10.64898/2026.05.13.725064

**Authors:** Samuel Ehrlich, Alexandra D. VandeLoo, Benjamin Magondu, Athena Chien, Sapna Sinha, Edward S. Boyden, Craig R. Forest

**Author notes:** Contributing authors.

## Abstract

A central challenge in opsin engineering is identifying mutations that reliably improve desired functional properties, a task made difficult by the enormous mutation space and limited throughput of electrophysiological screening. Improving opsin properties such as photocurrent amplitude and light sensitivity have the potential to broaden the use of opsins to low-light and deep-tissue applications. With this goal, we applied zero-shot protein language models (ESM-1b/1v) to recommend ChrimsonR mutations and experimentally validated all 17 of these variants using whole-cell patch clamp electrophysiology (n=6 cells per mutation). Despite many mutations reducing function, protein language models identified both known functional residues and unconventional substitutions that produced large functional gains and synergized with K176R to improve kinetics. Two mutations, E300G and E300P, increased sustained photocurrents from 66 pA (control) to 305 pA and 255 pA at 635 nm, reduced EC_50_ at 575 nm from 0.19 mW to 0.07 mW, and altered kinetics (*τ*_off_ increased from 0.06 s up to 0.40 s). Our results suggest that protein language models, even without task-specific training, can be used alongside electrophysiological measurements as a strategy for screening opsins for enhanced photocurrent.

## 1 Introduction

Opsins are a family of light-sensitive ion channels and pumps that have revolutionized neuroscience by enabling millisecond-scale, light-controlled modulation of neuronal activity [1, 2]. Their utility derives from key biophysical properties, including photocurrent amplitude, spectral sensitivity, and channel kinetics [3]. The engineering of opsins with enhanced or novel properties has advanced the field by providing tools with increased light sensitivity, altered spectra, or improved kinetics, achieved through rational design [4–8], exploration of natural diversity [9, 10], directed evolution [11], and more recently, supervised machine learning approaches [12, 13]. Despite these advances, there remain major unmet needs, including opsins with high photocurrents under low-light conditions, red-shifted spectra for deep tissue activation, and reduced immunogenicity for long-term clinical use.

A central challenge in opsin engineering is the vastness of the sequence space. For a protein the size of ChrimsonR (∼ 300 amino acids in the transmembrane domain), the number of possible single-site substitutions alone exceeds 19^300^ possible variants. Testing even a small fraction of these variants with patch clamp electrophysiology—the gold standard for ion channel function—is infeasible due to low throughput [14]. While innovations such as pipette cleaning [15] and automation [16–19] have increased efficiency, systematic exploration of the opsin sequence space is still beyond reach.

Protein language models (pLMs) offer a promising strategy to address this challenge. Trained on millions of natural protein sequences, these models capture statistical patterns that reflect evolutionary constraints and structural preferences [20, 21]. Remarkably, pLMs can predict the plausibility of mutations in a “zero-shot” setting—that is, without requiring task-specific training data. Hie et. al. demonstrated that such models could design novel antibodies with desirable properties directly from sequence likelihoods, bypassing traditional iterative screening [22]. This zero-shot paradigm is particularly attractive for opsins, where functional measurements are difficult and expensive to obtain, making them an ideal test case for leveraging pLMs to guide protein engineering.

Here, we applied a zero-shot likelihood scoring framework to three widely used opsins: ChrimsonR, ChR2, and C1C2. The model identified both conservative substitutions and unusual variants, including proline and glycine replacements within transmembrane *α*-helices, which are generally destabilizing but were selected as highly plausible. Using a recently developed robot for high throughput single-cell patch clamp electrophysiology [23], we systematically validated ChrimsonR variants. While most substitutions reduced or abolished photocurrents, two mutations at residue E300 (E300G and E300P) produced significantly enhanced sustained currents while slowing channel closing. Moreover, combining these amplitude-enhancing variants with the kinetic-accelerating mutation K176R generated channels with both larger currents and faster recovery, demonstrating the potential for synergistic engineering.

Together, these results highlight the utility of protein language models for navigating the immense opsin sequence space. By narrowing from billions of possible substitutions to a tractable set of candidates—including both previously known functional sites and unconventional variants—pLMs provide a powerful complement to experimental methods. When coupled with high-resolution functional assays such as patch clamp electrophysiology, this approach can accelerate the discovery of optogenetic tools with improved properties for neuroscience and beyond.

## 2 Results

### 2.1 Zero-shot predictions from protein language models across three opsins

In this study, a medley of six pre-trained models from the Evolutionary Scale Modeling (ESM) family were used developed by Meta. These were five versions of ESM-1b and one version of ESM-1v. The ESM-1b models were trained on UniRef50, a clustered database containing approximately 66 million sequence clusters, while ESM-1v was trained on UniRef90, containing roughly 208 million clusters by Meta. Of those, the UniRef50 database included 2 sequences annotated as “channelopsin” and 5 sequences annotated as “channelrhodopsin,” whereas UniRef90 contained 14 and 31 sequences for the same terms, respectively.

Modeling was performed following the zero-shot framework described by Hie et al. [22], in which models are trained on large protein sequence databases and can estimate the likelihood of individual residues or substitutions without task-specific fine-tuning. We applied this framework to ChrimsonR, ChR2, and C1C2, chosen for their common use and extensive engineering efforts. The models only output single-site substitutions by their evolutionary plausibility.

For ChrimsonR, ChR2, and C1C2 17, 27, and 24 mutations were predicted respectively. For all the opsins, the mutations spanned multiple transmembrane helices and loop regions (ChrimsonR results shown in Figure 1, others shown in Supplementary Table 1). Ten of the substitutions were chemically conservative (e.g., isoleucine to valine, phenylalanine to tryptophan) and 59 of the substitutions were non-conservative. To our knowledge, two predicted mutations have been described previously in the literature [6, 11] that were confirmed not to be present in the training dataset. The ChR2 mutation E90A has been found to increase photocurrent amplitude [11], and mutation C128T produces a step-opsin [8]. Additionally, previously identified important residues were chosen for substitution by the model, including the ChrimsonR residues E132 [24], K176 [9], and E300 [24].

**Fig. 1:**
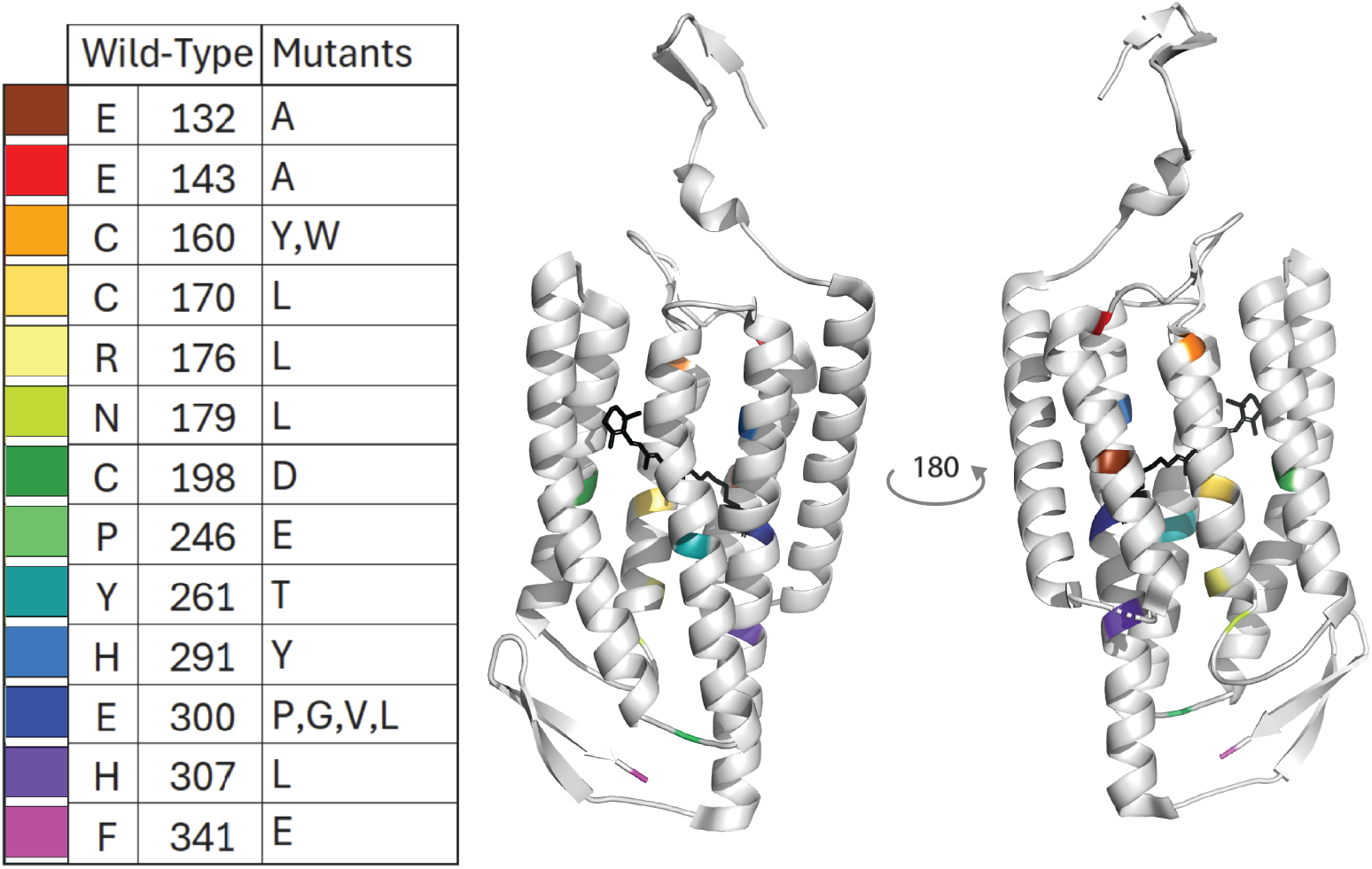
Color-coordinated list of the single amino acid ChrimsonR substitutions recommended by protein language model, with corresponding locations shown in the crystal structure(PDB: 5ZIH) from two angles.

Interestingly, the model also proposed mutations with unconventional substitutions. In particular, the substitutions E300P and E300G in ChrimsonR and N258P and N258G in ChR2 are notable because they introduce either a helix-breaking proline or a highly flexible glycine residue into the middle of a predicted transmembrane *α*-helix. In the following sections we chose to test the all the model recommended ChrimsonR mutants as there were more mutations that have not previously been discovered relative to ChR2 and C1C2.

### 2.2 Spectral Photocurrent responses of ChrimsonR mutations

All ChrimsonR mutants were tested for functional activity under whole-cell patch clamp in human embryonic kidney (HEK) cells with six different wavelengths. Averaged data revealed that 11 mutations resulted in either reduced photocurrent amplitude or complete loss of function across all tested wavelengths. Compared to the control, four substitutions (E132A, H307L, F341E, and E300V) maintained photocurrent, while two others, E300G and E300P, significantly increased sustained photocurrent amplitude to 305 pA and 255 pA respectively from the 66 pA control response with 635 nm light stimulation (Figure 2A, B).

**Fig. 2:**
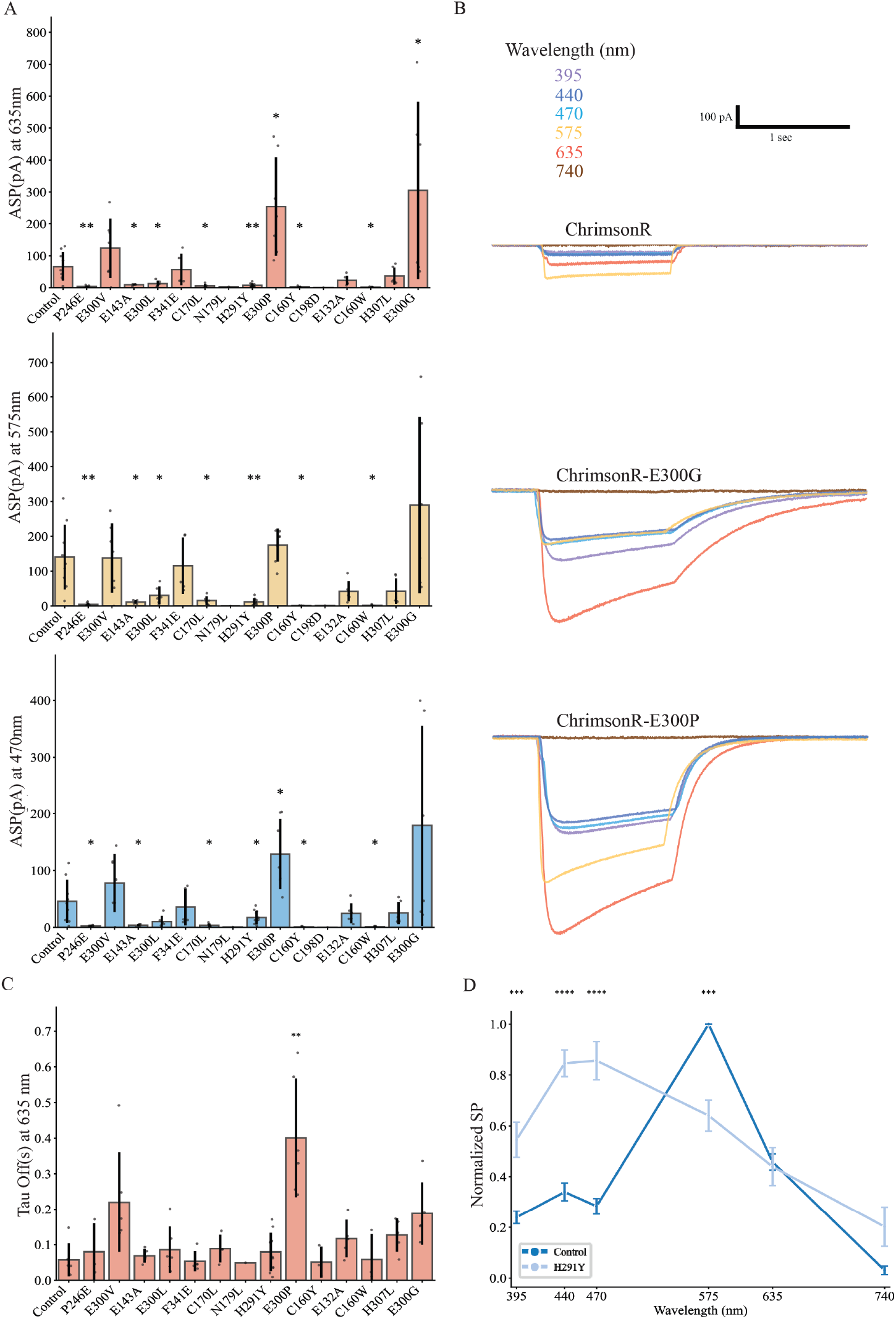
Photocurrent responses of protein language model recommended ChrimsonR mutations to light stimulation. **(A)** Average sustained photocurrent at 635 nm (red bars), 575 nm (yellow bars), and 470 nm (blue bars) across all measured mutations, showing that the mutations E300P and E300G produced significantly higher photocurrent relative to the unmutated control (255 pA, 305 pA, and 66 pA respectively).**(B)** Average sustained photocurrent traces across all measured wavelengths (colored according to legend) for E300G, E300P, and unmutated ChrimsonR, showing the fold increase in photocurrent for E300G and E300P. **(C)** Kinetic responses of mutants varied under the screening stimulation profile and were significantly different only for the E300P mutation. **(D)** Normalized sustained photocurrent as a function of excitation wavelength, showing a blue-shifted spectral response for mutation H291Y; all other mutation spectra overlap with WT, with no spectral shift (see appendices). (for all comparisons * signifies p < 0.05, Student’s t-test with Benjamini–Hochberg correction for multiple comparisons. n=6 cells for each wavelength)

One mutant displayed altered spectral sensitivity. The tyrosine substitution of the highly conserved residue [24] H291 reduced overall photocurrent to less than 20 pA for all wavelengths tested but produced a relative blue shift in wavelength dependence, with a peak response seen at 470 nm light stimulation (Figure 2C). This is the first reporting of this observation.

To quantify channel closing kinetics, we calculated the *τ*_off_ of photocurrent decay following light termination. E300 substitutions significantly prolonged *τ*_off_ relative to ChrimsonR and other mutants (control: 0.06 s, E300V: 0.22 s, E300G: 0.19 s, E300P:

0.40 s), consistent with their increased sustained photocurrent amplitudes (Figure 2D). To ensure that any measured differences in the electrophysiological response was due to changes in the channel itself and not due to expression, we measured the fluorescent expression of each mutant by quantifying integrated density. Fifteen out of seventeen mutants showed detectable fluorescence, and there was no significant difference in expression level between mutants (Supplementary Figure 1). Cells that were transfected with the remaining two mutants were not healthy enough to patch (data not shown).

### 2.3 E300P and G mutations increased sensitivity

We next investigated how the E300 substitutions affected light sensitivity by performing intensity sweeps at both 575 nm and 635 nm. At 575 nm, both E300P and E300G exhibited significantly lower EC_50_ (power required to yield half the maximum photocurrent, 0.07 mW for both relative to 0.19 mW for the control) values compared to ChrimsonR, indicating enhanced sensitivity to light. Importantly, for the 575 nm samples, maximum photocurrent amplitudes at higher photon fluxes were not different from the control. This suggests that the primary effect of these mutations is to reach saturation with less light intensity (Figure 3A, B, C). Despite higher measured photocurrent, at 635 nm stimulation there was no measurable difference in EC_50_ (Supplementary Figure 2).

**Fig. 3:**
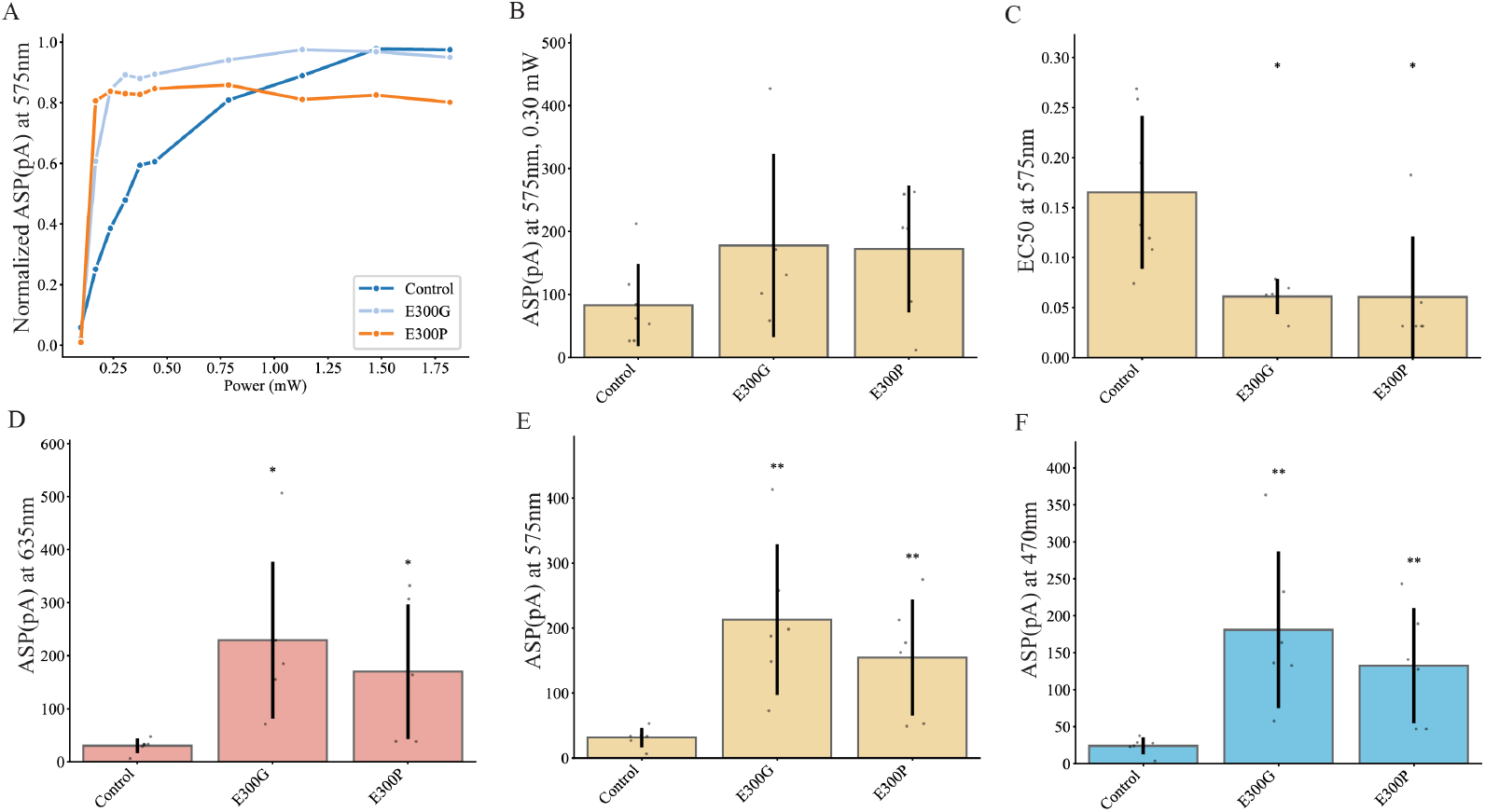
Photocurrent responses of E300P and E300G mutations to light stimulation of varying single wavelength intensity stimulation or reduced intensity multiwavelength stimulation. (A) Normalized sustained photocurrent at 575 nm for the ChrimsonR control compared to the pLM suggested E300 variants shows that P and G responses saturate at 0.25 mW compared to 1.5 mW in the control. (B) The average sustained photocurrent amplitude at 0.30 mW from panel A. There was no significant difference in amplitude at any intensity. (C) The EC_50_ in mW of intensity sweeps with 575 light showed that the photocurrent amplitude reached 50% of it maximum at nearly half the power for the E300G and P mutations compared to the control (0.07 vs 0.19 mW). (D,E,F) The average sustained photocurrent of the E300P and G mutations compared to the controls at 635, 575, and 470nm respectively shows that E300P and G significantly increased the photocurrent response at all wavelengths. (* indicates p-value < 0.05, calculated by student t-test adjusted by the Benjamini-Hochberg procedure to account for multiple comparisons)

To assess whether this difference in EC_50_ response at 575 nm is due to saturation of the channel, we used the same stimulation protocol as described in the previous section at reduced power. We first determined the power required to nearly saturate the photocurrent response of E300G and E300P with 635 nm light (80% of maximum response). The other five wavelengths were scaled to have equivalent photon flux. Under these lower light conditions, both E300P and E300G produced significantly larger sustained photocurrents (170 pA and 229 pA respectively) compared to ChrimsonR (30 pA) (Figure 3D) with no difference in the spectral response, as expected (Supplementary Figure 3).

### 2.4 Mutations that increase closing kinetic speed and photocurrent amplitude work synergistically

To assess whether combining mutations with distinct effects on channel properties could yield additive or synergistic outcomes, we introduced the amplitude-enhancing variants E300G and E300P into Chrimson Wild-Type plasmids which lack the kinetic-accelerating mutation K176R. To accurately quantify both peak photocurrent amplitude and closing kinetics, we employed a modified stimulation protocol using brief 5 ms light pulses during 635 nm intensity sweeps. Representative traces (Figure 4A) demonstrate that inclusion of the K176R substitution consistently accelerated channel closing across all variants. Analysis of closing kinetics revealed a robust effect: *τ*_off_ was significantly reduced in all channels carrying the K176R substitution compared to their R176 counterparts with a decrease in *τ*_off_ of 0.35 s and 0.65 s for E300G and E300P respectively. (Figure 4B). As expected, intensity–response curves (Figure 4C) showed that the combined mutations did not alter light sensitivity. We also found no significant difference in peak current amplitude under these conditions (Supplementary Figure 4).

**Fig. 4:**
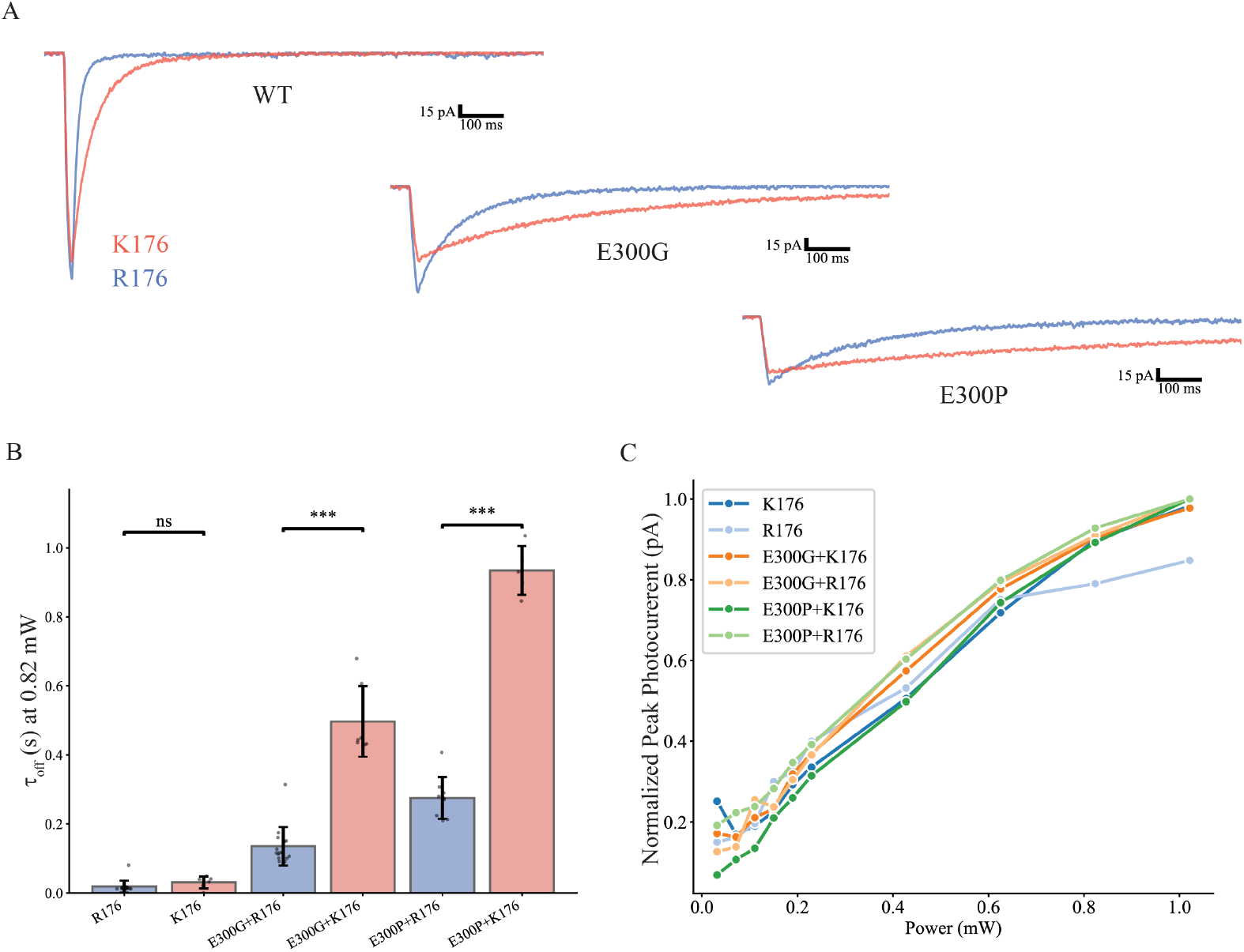
Graphs comparing the mutations E300G and E300P in wild-type Chrimson and ChrimsonR (which has the K176R kinetics increasing mutation and was used as the control throughout this paper). (A) Comparison between the R176 (blue) and K176 (red) overlaid raw photocurrent responses show that mutations for increasing kinetic speed (K176R) in the WT Chrimson also increase the kinetic speed with mutations E300G and E300P present. (B) *τ*_off_ measured with 635nm light at 0.82 mW compared across the mutants and control with and without the K176R mutation, shows that R176 results in significantly faster(decreased tau off) response in all cases. (C) Normalized peak photocurrent of ChrimsonR and the various mutations across a power sweep. (*** indicates p-value < 0.001, ** indicates p-value < 0.01, * indicates p-value < 0.05, calculated by student t-test adjusted by the Benjamini-Hochberg procedure to account for multiple comparisons)

## 3 Discussion

We explored the application of protein language models to guide opsin engineering and experimentally validated their predictions through patch-clamp electrophysiology. These models were not specifically trained on channelrhodopsins and were solely trained amino acid sequences, with no knowledge of protein structure or function. We first validated the models’ by identifing reccomended substitutions in Chrim-sonR, ChR2, and C1C2 that were ranked as evolutionarily plausible by the model and comparing them to known mutations in literature. The full list can be found in Supplementary Table 1. Encouragingly, many residues previously implicated in opsin function were recommended. Specifically in ChrimsonR the E132, K176, and E300 locations were recommended, highlighting the models’ ability to recommend unconventional yet promising channelrhodopsin mutations with enhanced photocurrents despite not being fine-tuned on opsin sequence, structure or function. Interestingly, across all three opsins, while several substitutions were chemically conservative, the majority were non-conservative, highlighting the model’s ability to suggest unusual variants that might not be considered through traditional rational design (Figure 1). These encouraging results led us to probe further experimentally into the function of the suggested ChrimsonR mutations.

Our electrophysiological characterization of ChrimsonR variants resulted in the discovery of three particularly interesting single amino acid mutations. E300P and E300G increased sustained photocurrent amplitude with 635 nm light stimulation by 189 pA and 239 pA respectively. These findings extend prior work implicating E300 as a functionally important site [24] and further reveal that substitutions of proline and glycine in a helix, although typically considered destabilizing, can nonetheless enhance photocurrent amplitude. H291Y blue-shifted the wavelength response from 575 nm to 470 nm with our six wavelength stimulation protocol. These results illustrate that single mutations suggested by the pLM can significantly alter biophysical properties.

We found the observance of proline and glycine residues in helices to be relatively absent in the literature [25], making the strong effects of the intrahelical mutations at E300 particularly interesting. We hypothesized that, because the pLMs were trained on protein sequences from the UniRef50 and UniRef90 databases, they would recommend mutations based on patterns previously observed in protein sequences. However, we found less than 10 matching seven-amino-acid sequences in databases (more than 4 million sequences in each UniRef50 and UniRef90 databases). Also, of the matches with a solved secondary structure, the sequence was never located in an alpha-helix. Therefore, the pLM suggestion is also likely not from inferred secondary structure, unless these mutations caused the helix to fully destabilize while somehow still allowing the protein to fold and function. To explore this further, future work could solve the structure of this mutated ChrimsonR to identify whether structural changes caused by the glycine and proline mutations were responsible for the increased photocurrent response.

While surprising that the E300P, G mutations so greatly increased photocurrent, the associated slowing of the closing kinetics has been observed by others [24]. To account for the contribution of channel closing kinetics to steady-state photocurrent, we compared a kinetically normalized metric defined as sustained photocurrent divided by *τ*_off_. Using reported steady-state photocurrents for representative channel-rhodopsins, fast variants such as ChR2 exhibit high normalized values ( 40,000 pA/s [1]), reflecting rapid channel closing and moderate steady-state current. Red-shifted variants such as ReaChR occupy an intermediate regime ( 3000 pA/s [4]), balancing slower kinetics with larger sustained currents. Within this framework, ChrimsonR has a normalized photocurrent of 1000 pA/s with E300G exceeding the control with 1600 pA/s, indicating that its increased sustained photocurrent is likely not solely explained by slower channel closing. In contrast, E300P exhibits a reduced normalized value of 600 pA/s, consistent with a dominant contribution from prolonged open-state lifetime. It should be noted that these comparisons are approximate and future work could evaluate the E300 mutated ChrimsonR to existing optogentic tools under equivalent conditions.

However, we found that combining the kinetic-accelerating mutation K176R with amplitude-enhancing substitutions at E300 yielded channels that recovered more rapidly while maintaining higher sustained photocurrents (Figure 4). This demonstrates that rationally combining mutations with complementary effects can produce synergistic variants, enabling the simultaneous optimization of multiple channel properties.

In addition to the beneficial substitutions, the model proposed several mutations that did not translate into qualitatively novel or exotic channel properties. Also, none of the mutations seemed to increase the speed of the channels, or narrow their response spectra, both of which would be highly desirable changes. These results highlight an important limitation to the current model: while protein language models can identify beneficial substitutions, they do not explicitly optimize for specific, desirable functional properties, such as ultrafast kinetics. Due to their training on known protein sequences, pLMs may be more effective at nominating stable, evolutionarily-tolerant substitutions than at discovering rare mutations that confer exciting new optogenetic capabilities.

Our results suggest that protein language models can serve as powerful tools for recommending unconventional yet promising channelrhodopsin mutations with enhanced photocurrents. The zero-shot scoring framework reduced the search space to a tractable set of candidates, including both previously validated variants and novel substitutions with strong functional effects. Importantly, these predictions require no prior knowledge of opsin function or retraining on domain-specific data, enabling generalizable applications across other membrane proteins. Recent work has begun to demonstrate the utility of combining pLMs with large experimental opsin datasets to more directly learn sequence–function relationships [26], suggesting that richer training data could further enhance the predictive capacity of these models. As such datasets continue to grow, future design frameworks that integrate pLM priors with explicit functional objectives may unlock more ambitious engineering goals, including tailored spectral sensitivity, accelerated kinetics, or improved membrane trafficking.

## 4 Methods

### 4.1 Cell culture

HEK293FT cells (Invitrogen) were selected as the expression host for several reasons: (i) they are highly amenable to lipofectamine transfection; (ii) they are widely used for robust expression of exogenous proteins, making them ideal for optogenetic tool characterization; (iii) they are reported to have among the lowest rates of exogenous DNA-induced mutation among commonly used mammalian cell lines; (iv) they are easy to grow and maintain, and widely adopted for tool development and screening.

Cells were cultured at 37°C with 5% CO_2_ in DMEM (Gibco) supplemented with 10% heat-inactivated fetal bovine serum (FBS, Corning), and 1% sodium pyruvate (Gibco). Cells were maintained between 10% and 70% confluence to preserve exponential growth. Cells were seeded onto coverslips (0.15mm thick, 12mm in diameter, Electron Microscopy Systems) coated in Matrigel (2% growth factor–reduced Matrigel, Corning) diluted in DMEM for 1h at 37 °C.

### 4.2 Cell transfection

Mutant plasmids were ordered from Azenta Life Sciences, plasmid map for ChrimsonR experiments in Supplementary Figure 6. Approximately 70,000 cells were seeded onto the coverslips and transfected the following day using 1µl of Lipofectamine 2000 (Invitrogen) and 150 ng of DNA. After one day of incubation the cells were trypsinized and placed on new coverslips at a lower density.

### 4.3 Zero shot protein language model prediction

Protein language models were used to prioritize amino acid substitutions for opsin engineering based on evolutionary plausibility. Modeling was performed following the zero-shot framework described by Hie et al. [22], in which models are trained on large protein sequence databases and can estimate the likelihood of individual residues or substitutions without task-specific fine-tuning. The code developed by Hie et. al. was downloaded from github (https://github.com/brianhie/efficient-evolution) and used without alteration. A protein sequence is simply entered as an input and a list of amino acid substitutions is returned along with the number of models recommending the substitution (out of 6).

### 4.4 Electrophysiology instrumentation

All patch clamp recordings were conducted using an Axopatch 200B amplifier (Molecular Devices). The analog voltage output from the amplifier was digitized and recorded using a National Instruments data acquisition device (NI USB-6210, National Instruments), which provided synchronized analog-to-digital conversion and timing control for stimulus delivery and data acquisition.

The patch pipette was mounted on a three-axis micromanipulator (Sensapex), which enabled fine automated positioning and alignment for seal formation and whole-cell configuration. Fluorescence excitation for identifying opsin-expressing cells was provided by a mercury arc lamp (X-Cite 120, Excelitas Technologies), with standard filter sets for tdTomato and eGFP (Thermo Fischer Scientific).

Optogenetic stimulation was delivered through the patch pipette using an LED light engine (Lumencor Spectra X), which provided illumination across multiple wavelengths. Light was coupled into the back of the patch pipette through an optical fiber inserted into a special purpose pipette holder (Optopatcher) [27], enabling precisely colocalized stimulation and recording.

Pressure control for the patch pipette was maintained using a custom-built multi-channel pressure control box that delivered computer-regulated positive and negative pressures for cell access and seal formation as previously described [17]. A single pipette was reused unless it failed due to clogging, breaking, or repeated failure to form a gigaseal.

### 4.5 Electrophysiology recording

The external (bath) solution was Tyrode’s solution, containing (in mM): 125 NaCl, 2 KCl, 3 CaCl_2_, 1 MgCl_2_, 10 HEPES, and 30 glucose (pH adjusted to 7.3 with NaOH, 305 mOsm). The internal pipette solution contained (in mM): 125 K-gluconate, 8 NaCl, 0.1 CaCl_2_, 0.6 MgCl_2_, 1 EGTA, 10 HEPES, 4 Mg-ATP, and 0.4 Na-GTP (pH adjusted to 7.3 with KOH, 295–300 mOsm).

For recording, we manually selected cells exhibiting the following criteria: (1) non-zero fluorescence visible at set exposure (1 s on our camera) and (2) physically isolated from neighboring cells. Whole-cell patch-clamp recordings were performed on isolated HEK293FT cells to minimize space-clamp artifacts and ensure accurate measurements of photocurrents. All recordings were conducted at room temperature. All data was collected with a 10 kHz lowpass Bessel filter applied.

Patched cells which did not meet the following patch quality control threshholds were rejected from analysis: access resistance <35MΩ and holding current within *±*100pA. Typical membrane resistance ranged from 200MΩ to 2GΩ, and pipette resistance was 3–9MΩ. Patch pipettes were pulled from borosilicate glass with a 1.5mm outside diameter and a 0.86mm inside diameter (Warner Instruments).

Each cell underwent a brief voltage-clamp test step of +10mV from holding potential to assess membrane capacitance and seal quality. This was followed by a current-clamp protocol consisting of three consecutive current injections (-10, 0, and +10pA) to confirm membrane responsiveness and resting potential stability. After patching, pipettes were cleaned as previously described using tergazyme (Alconox) [15].

#### 4.5.1 Optical stimulation

To determine the optical stimulation protocol we first found the minimum illumination power needed to produce the maximum photocurrent amplitude of the ChrimsonR sequence and introduce a safety factor in order to reduce the risk of photobleaching during experiments. This calibration was done by performing intensity sweeps to determine the optimal illumination power at the 635 nm wavelength as previously reported [23]. We chose to set the illumination power at approximately 75% of the power at maximum sustained photocurrent amplitude for the experiments herein (0.842 mW as measured at the end of the optical fiber). The other excitation wavelengths were calibrated to match the photon flux of the chosen power for the 635 nm light.

The light protocol consisted of three randomized sweeps per cell, each containing six discrete wavelengths: 395, 440, 470, 575, 635, and 740 nm. For each sweep, cells were held in darkness for 15 s, stimulated for 1 s with a single wavelength, and returned to darkness for 14 s, totaling 90 s per full stimlution series while being held by voltage clamp at -50 mV. This holding potential was selected to increase the amplitude of the photocurrent response which increased the signal to noise ratio of the recorded response. Wavelengths were randomized across trials to control for order effects. All light intensities were pre-calibrated for consistent photon flux delivery across wavelengths.

In addition, further protocols were employed to probe response at lower power. For these measurements, illumination power was reduced to 0.131 mW at 635 nm of the calibrated intensity (with all other wavelengths set to equal photon flux) to generate submaximal responses, minimizing channel saturation effects. Additionally, the cells were stimulated a single time per wavelength to further reduce saturation effects. To probe kinetics and peak currents, cells were also subjected to intensity sweeps with 5 ms light pulses at 635 nm. Short-pulse stimulation allowed for improved resolution of channel activation and deactivation kinetics, while also enabling precise measurement of peak photocurrent amplitude under rapid excitation. Similar to the low power stimulation, cells were stimulated only once per intensity to minimize saturation effects.

### 4.6 Data analysis

Electrophysiology data was analyzed using custom Python scripts. All data was filtered with a digital 3kHz Bessel filter after data collection. Sustained photocurrents were defined as the average of the photocurrent from the baseline holding current during the last 50ms of light stimulation. Cells which exhibited sustained photocurrents greater than 15pA were included in spectral response analysis. Peak currents were defined as the maximum deviation from baseline current during light stimulation. *τ*_off_ was calculated by fitting the response to a monoexponential equation at the end of light stimulation.

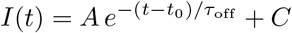

EC_50_ was calculated by fitting the measured sustained photocurrents across multiple intensities of light to the Hill equation.

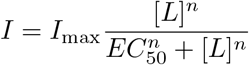

Fluorescent expression was determined using ImageJ, cells were manually traced and fluorescent parameters such integrated density were automatically generated by ImageJ. Error bars indicate the standard deviation (SD) in all figures. Statistical significance was determined by a conducting a student t-test between control and experimental conditions. To account for error due to multiple comparisons the Benjamini-Hochberg procedure was applied.

## 5 Acknowledgements

CRF acknowledges the NIH BRAIN Initiative Grant (NEI and NIMH 1-U01-MH106027-01), NIH R01NS102727, NIH Single Cell Grant 1 R01 EY023173, NIH R01DA029639 and NIH RF1AG079269, support from Georgia Tech through the Institute for Bioengineering and Biosciences, Invention Studio, and the George W. Woodruff School of Mechanical Engineering. S.S. acknowledges the Schmidt Science Fellows for their generous support through the postdoctoral fellowship. ESB acknowledges Lisa Yang, HHMI, NIH 1R01MH123977, NIH R01MH122971, and NIH R01DA029639. ChatGPT was used in the editing of this work.

## 6 Author conritbutions

**SE**: Conceptualization, Data curation, Formal analysis, Investigation, Methodology, Software, Validation, Visualization, Writing – original draft, Writing – review and editing. **ADV**: Visualization, Writing – review and editing. **BM**: Methodology. **AC**: Writing – review and editing. **SS**: Writing – review and editing. **ESB**: Conceptualization, Funding acquisition, Methodology, Supervision, Writing – review and editing. **CRF**: Conceptualization, Data curation, Formal analysis, Funding acquisition, Methodology, Supervision, Writing – original draft, Writing – review and editing.

## Declarations

ESB is an inventor on several patents related to optogenetics.

## References

[1] Boyden, E.S., Zhang, F., Bamberg, E., Nagel, G., Deisseroth, K.: Millisecond-timescale, genetically targeted optical control of neural activity. Nature Neuroscience 8, 1263–1268 (2005) 10.1038/nn1525

[2] Deisseroth, K., Hegemann, P.: The form and function of channelrhodopsin. Science 357, 5544 (2017) https://doi.org/10.1126/science.aan5544. doi: 10.1126/science.aan5544

[3] Piatkevich, K.D., Boyden, E.S.: Optogenetic control of neural activity: the biophysics of microbial rhodopsins in neuroscience. Quarterly Reviews of Biophysics 57, 1 (2024)

[4] Lin, J.Y., Knutsen, P.M., Muller, A., Kleinfeld, D., Tsien, R.Y.: Reachr: a red-shifted variant of channelrhodopsin enables deep transcranial optogenetic excitation. Nature Neuroscience 16, 1499–1508 (2013) 10.1038/nn.3502

[5] Nagel, G., Brauner, M., Liewald, J.F., Adeishvili, N., Bamberg, E., Gottschalk, A.: Light activation of channelrhodopsin-2 in excitable cells of caenorhabditis elegans triggers rapid behavioral responses. Current biology 15, 2279–2284 (2005)

[6] Gunaydin, L.A., Yizhar, O., Berndt, A., Sohal, V.S., Deisseroth, K., Hegemann, P.: Ultrafast optogenetic control. Nature Neuroscience 13, 387–392 (2010)10.1038/nn.2495

[7] Gong, X., Mendoza-Halliday, D., Ting, J.T., Kaiser, T., Sun, X., Bastos, A.M., Wimmer, R.D., Guo, B., Chen, Q., Zhou, Y., Pruner, M., Wu, C.W.-H., Park, D., Deisseroth, K., Barak, B., Boyden, E.S., Miller, E.K., Halassa, M.M., Fu, Z., Bi, G., Desimone, R., Feng, G.: An ultra-sensitive step-function opsin for minimally invasive optogenetic stimulation in mice and macaques. Neuron 107, 38–518 (2020) https://doi.org/10.1016/j.neuron.2020.03.032. doi: 10.1016/j.neuron.2020.03.032

[8] Berndt, A., Yizhar, O., Gunaydin, L.A., Hegemann, P., Deisseroth, K.: Bi-stable neural state switches. Nature neuroscience 12, 229–234 (2009)

[9] Klapoetke, N.C., Murata, Y., Kim, S.S., Pulver, S.R., Birdsey-Benson, A., Cho, Y.K., Morimoto, T.K., Chuong, A.S., Carpenter, E.J., Tian, Z., Wang, J., Xie, Y., Yan, Z., Zhang, Y., Chow, B.Y., Surek, B., Melkonian, M., Jayaraman, V., Constantine-Paton, M., Wong, G.K.-S., Boyden, E.S.: Independent optical excitation of distinct neural populations. Nature Methods 11, 338–346 (2014) 10.1038/nmeth.2836

[10] Marshel, J.H., Kim, Y.S., Machado, T.A., Quirin, S., Benson, B., Kadmon, J., Raja, C., Chibukhchyan, A., Ramakrishnan, C., Inoue, M.: Cortical layer–specific critical dynamics triggering perception. Science 365, 5202 (2019)

[11] Cho, Y.K., Park, D., Yang, A., Chen, F., Chuong, A.S., Klapoetke, N.C., Boyden, E.S.: Multidimensional screening yields channelrhodopsin variants having improved photocurrent and order-of-magnitude reductions in calcium and proton currents. Journal of Biological Chemistry 294, 3806–3821 (2019)10.1074/jbc.RA118.006996. doi: 10.1074/jbc.RA118.006996

[12] Bedbrook, C.N., Yang, K.K., Rice, A.J., Gradinaru, V., Arnold, F.H.: Machine learning to design integral membrane channelrhodopsins for efficient eukaryotic expression and plasma membrane localization. PLoS computational biology 13, 1005786 (2017)

[13] Bedbrook, C.N., Yang, K.K., Robinson, J.E., Mackey, E.D., Gradinaru, V., Arnold, F.H.: Machine learning-guided channelrhodopsin engineering enables minimally invasive optogenetics. Nature Methods 16, 1176–1184 (2019)10.1038/s41592-019-0583-8

[14] Hamill, O.P., Marty, A., Neher, E., Sakmann, B., Sigworth, F.J.: Improved patch-clamp techniques for high-resolution current recording from cells and cell-free membrane patches. Pflügers Archiv 391, 85–100 (1981)10.1007/BF00656997

[15] Kolb, I., Stoy, W.A., Rousseau, E.B., Moody, O.A., Jenkins, A., Forest, C.R.: Cleaning patch-clamp pipettes for immediate reuse. Scientific Reports 6, 35001 (2016) 10.1038/srep35001

[16] Kodandaramaiah, S.B., Franzesi, G.T., Chow, B.Y., Boyden, E.S., Forest, C.R.: Automated whole-cell patch-clamp electrophysiology of neurons in vivo. Nature Methods 9, 585–587 (2012) 10.1038/nmeth.1993

[17] Kolb, I., Landry, C.R., Yip, M.C., Lewallen, C.F., Stoy, W.A., Lee, J., Felouzis, A., Yang, B., Boyden, E.S., Rozell, C.J., Forest, C.R.: Patcherbot: a single-cell electrophysiology robot for adherent cells and brain slices. Journal of Neural Engineering 16, 046003 (2019) 10.1088/1741-2552/ab1834

[18] Yip, M.C., Gonzalez, M.M., Valenta, C.R., Rowan, M.J.M., Forest, C.R.: Deep learning-based real-time detection of neurons in brain slices for in vitro physiology. Scientific reports 11, 6065 (2021)

[19] Gonzalez, M.M., Lewallen, C.F., Yip, M.C., Forest, C.R.: Machine learning-based pipette positional correction for automatic patch clamp in vitro. Eneuro 8 (2021)

[20] Lin, Z., Akin, H., Rao, R., Hie, B., Zhu, Z., Lu, W., Smetanin, N., Verkuil, R., Kabeli, O., Shmueli, Y., Santos Costa, A., Fazel-Zarandi, M., Sercu, T., Candido, S., Rives, A.: Evolutionary-scale prediction of atomic-level protein structure with a language model. Science 379 (2023) 10.1126/science.ade2574

[21] Hayes, T., Rao, R., Akin, H., Sofroniew, N.J., Oktay, D., Lin, Z., Verkuil, R., Tran, V.Q., Deaton, J., Wiggert, M.: Simulating 500 million years of evolution with a language model. Science 387, 850–858 (2025)

[22] Hie, B.L., Shanker, V.R., Xu, D., Bruun, T.U.J., Weidenbacher, P.A., Tang, S., Wu, W., Pak, J.E., Kim, P.S.: Efficient evolution of human antibodies from general protein language models. Nature biotechnology 42, 275–283 (2024)

[23] Ehrlich, S., VandeLoo, A.D., Badawy, M., Gonzalez, M.M., Stockslager, M., Yang, A., Sinha, S., Bracha, S., Park, D., Magondu, B., et al.: Screening channel-rhodopsins using robotic intracellular electrophysiology and single cell sequencing. Journal of Neuroscience Methods, 110663 (2025)

[24] Vierock, J., Grimm, C., Nitzan, N., Hegemann, P.: Molecular determinants of proton selectivity and gating in the red-light activated channelrhodopsin chrimson. Scientific Reports 7, 9928 (2017)

[25] Yun, R.H., Anderson, A., Hermans, J.: Proline in α-helix: Stability and conformation studied by dynamics simulation. Proteins: Structure, Function, and Bioinformatics 10, 219–228 (1991)

[26] Tose, A.J., Nava, A.A., McGrath, S.N., Mardinly, A.R., Naka, A.: Wachrs are excitatory opsins sensitive to indoor lighting. bioRxiv, 2025–2029 (2025)

[27] Katz, Y., Yizhar, O., Staiger, J., Lampl, I.: Optopatcher—an electrode holder for simultaneous intracellular patch-clamp recording and optical manipulation. Journal of neuroscience methods 214, 113–117 (2013)

