## Supplemental figures for "Mutation E300 recommended by protein language models gives ChrimsonR amplified photocurrent response"

Supplementary figures for “Protein language-models recommend unconventional yet promising channelrhodopsin mutations with enhanced photocurrents”

| Channelrhodopsin-2 | ChrimsonR | C1C2 |
|---|---|---|
| F16N | E132A | S1M |
| M68V | E143A | I44V |
| C79K | C160Y | M84V |
| E83L | C160W | W73A |
| E90L | C170L | C95F |
| E90A | R176L | E106A |
| M91I | N179L | K109Y |
| K93A | C198D | K109L |
| K93G | P246E | E117F |
| K93L | Y261T | E117A |
| E97Y | H291Y | E117L |
| C128T | E300P | C144T |
| N137L | E300G | N153L |
| C183V | E300V | S171A |
| N187Y | E300L | C199S |
| N187L | H307L | P220K |
| P204G | F341E | C224K |
| C208R | | C224L |
| F226Y | | F246W |
| F230W | | H265Y |
| H249Y | | N274L |
| N258G | | N274P |
| N258L | | G280L |
| N258P | | H281L |
| G264L | | |
| H265L | | |
| T313K | | |

*Supplementary Table 1. The combined list of predicted amino acid substitutions for three opsins output by the protein language model, Evolutionary Scale Modeling versions 1b and 1v, ordered by location in the protein based on their published sequences in the Protein DataBase (PDB).*

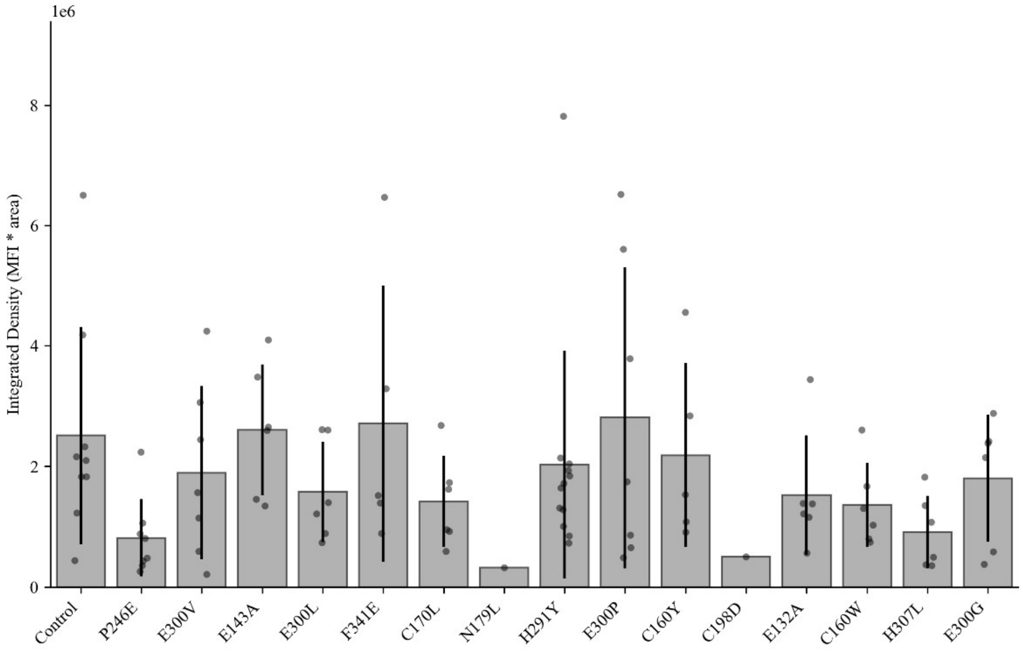


*Supplementary Figure 2. Fluorescence expression (calculated using mean fluorescence intensity metric in ImageJ) of cells expressing the PLM recommended mutations in ChrimsonR.*

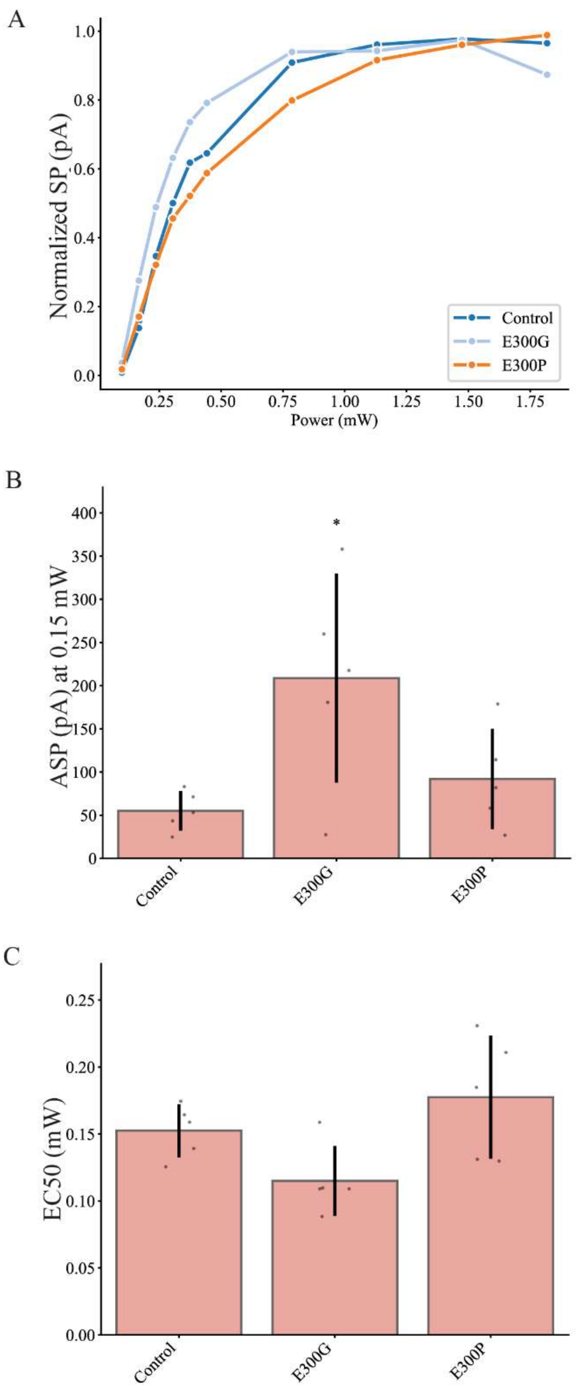


*Supplementary Figure 3. Normalized sustained photocurrent at 635 nm for the Chrimson R control compared to the 4 PLM suggested E300 variants. (B) Zoomed in representation of the $EC_{50}$ of intensity sweeps in mW with 635nm light showed that the photocurrent amplitude did not significantly differ from the control at this 635nm in contrast to the response at 575 nm. (C) The sustained photocurrent at low power (equivalent to $EC_{50}$) showed significantly higher photocurrent compared to the control in E300G, and significantly lower in E300L.*

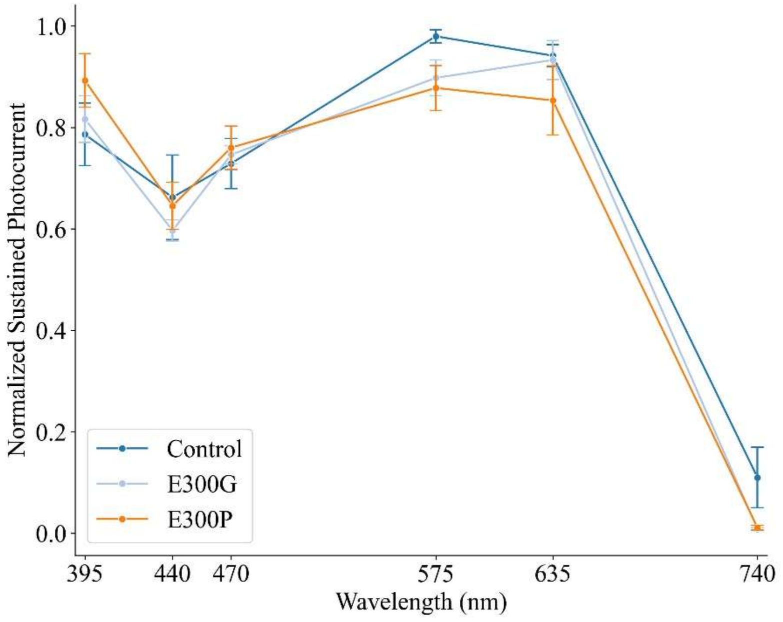


*Supplementary Figure 4. Normalized sustained photocurrent of ChrimsonR and E300 mutants using reduced intensity of light stimulation.*

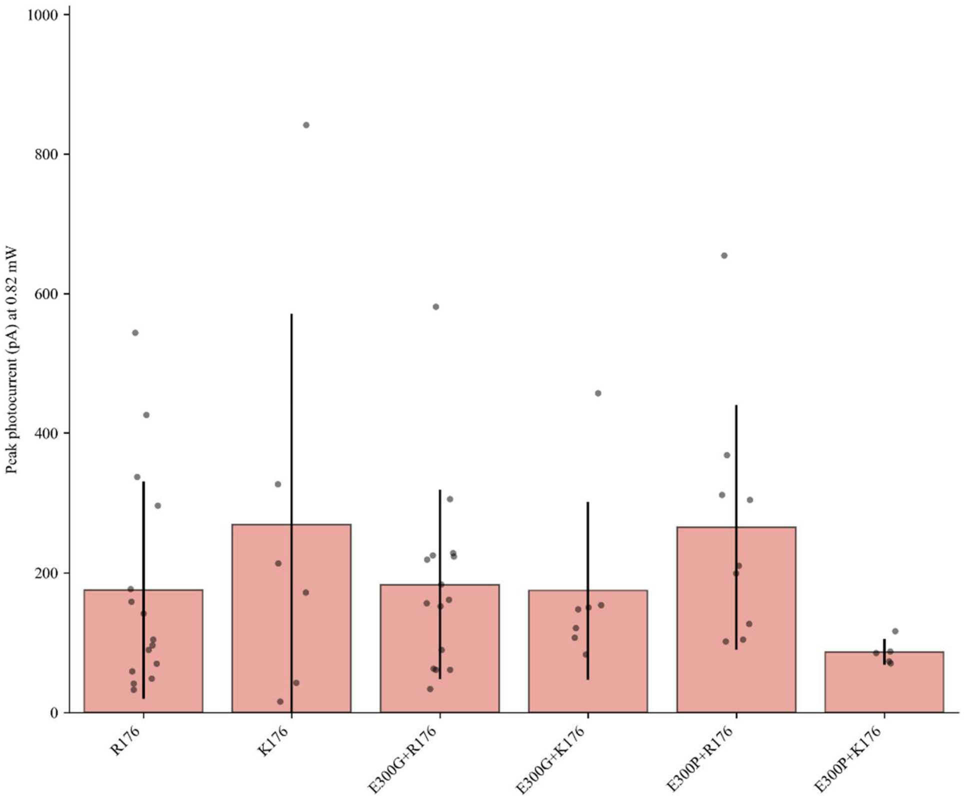


*Supplementary Figure 5. Peak photocurrents of ChrimsonR with and without the mutations as shown on the axis at the power indicated on the y-axis.*

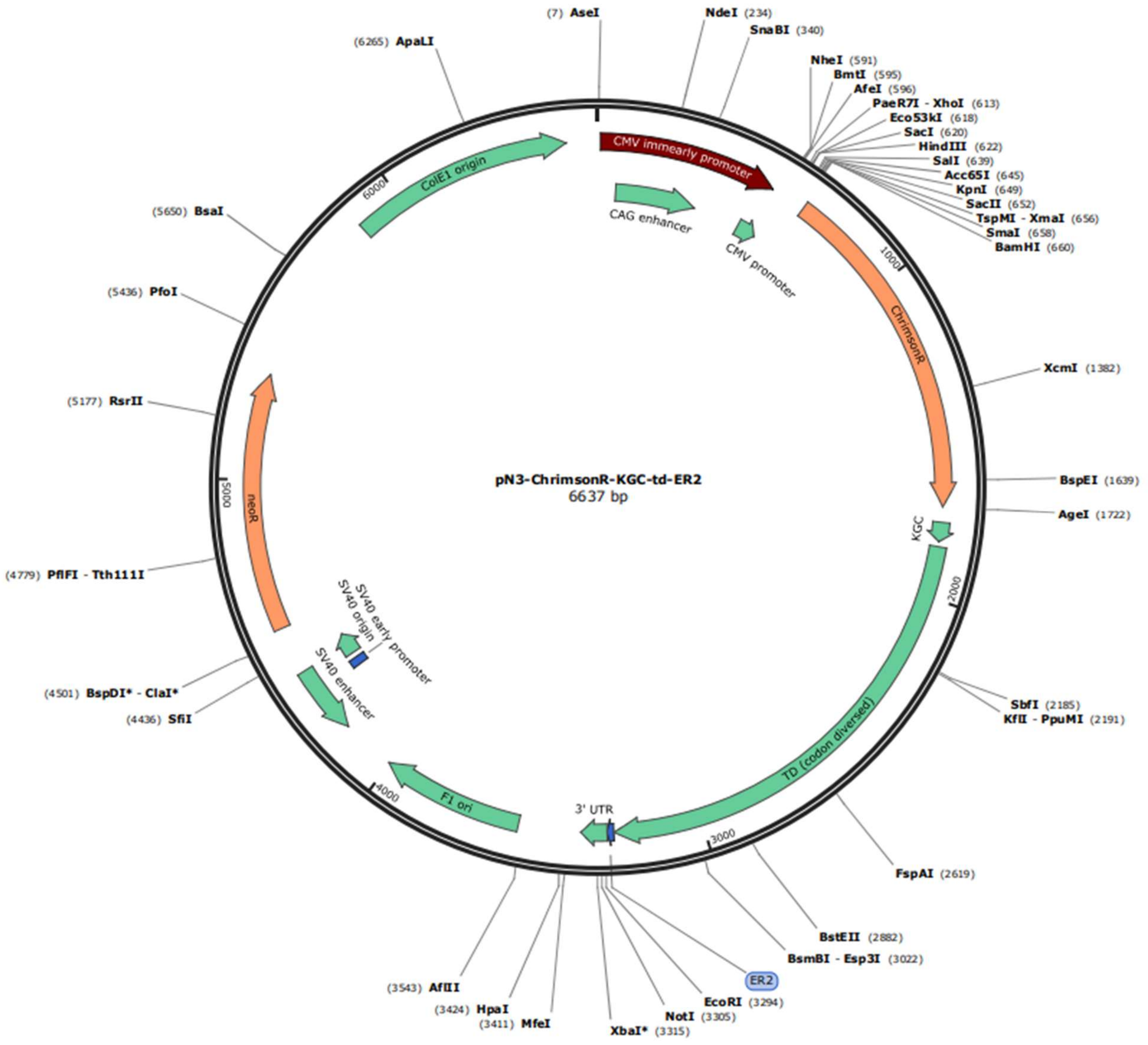


*Supplementary Figure 6. Representative map of plasmids used for transfection, containing ChrimsonR channel rhodopsin fused to tdtomato fluorophore. The so named pN3-ChrimsonR-KGC-td-ER2 plasmid was used as the control, and all PLM-suggested single amino-acid mutants were directly edited within this plasmid in order to maintain a controlled system.*
